# Evaluation of DNA extraction methods on individual helminth egg and larval stages for whole genome sequencing

**DOI:** 10.1101/616672

**Authors:** Stephen R. Doyle, Geetha Sankaranarayan, Fiona Allen, Duncan Berger, Pablo D. Jimenez Castro, James Bryant Collins, Thomas Crellen, María A. Duque-Correa, Peter Ellis, Tegegn G. Jaleta, Roz Laing, Kirsty Maitland, Catherine McCarthy, Tchonfienet Moundai, Ben Softley, Elizabeth Thiele, Philippe Tchindebet Ouakou, John Vianney Tushabe, Joanne P. Webster, Adam J. Weiss, James Lok, Eileen Devaney, Ray M. Kaplan, James A. Cotton, Matthew Berriman, Nancy Holroyd

**Author notes:** Correspondence: Stephen R. Doyle, Nancy Holroyd.

## Abstract

Whole genome sequencing is being rapidly applied to the study of helminth genomes, including *de novo* genome assembly, population genetics, and diagnostic applications. Although late-stage juvenile and adult parasites typically produce sufficient DNA for molecular analyses, these parasitic stages are almost always inaccessible in the live host; immature life stages found in the environment for which samples can be collected non-invasively offer a potential alternative, however, these samples are typically yield very low quantities of DNA, can be environmentally resistant, and are susceptible to contamination, often from bacterial or host DNA. Here, we have tested five low-input DNA extraction protocols together with a low-input sequencing library protocol to assess the feasibility of whole genome sequencing of individual immature helminth samples. These approaches do not use whole genome amplification, a common but costly approach to increase the yield of low input samples. We first tested individual parasites from two species spotted onto FTA cards - egg and L1 stages of *Haemonchus contortus* and miracidia of *Schistosoma mansoni* - before further testing on an additional six species - *Ancylostoma caninum, Ascaridia dissimilis, Dirofilaria immitis, Dracunculus medinensis, Strongyloides stercoralis, and Trichuris muris* - with an optimal protocol. Whole genome sequencing followed by analyses to determine the proportion of on- and off-target mapping revealed successful sample preparations for six of the eight species tested with variation between species, and within species but between life stages, described. These results demonstrate the feasibility of whole genome sequencing of individual parasites, and highlight a new avenue towards generating sensitive, specific, and information-rich data for the diagnosis and surveillance of helminths.

## Introduction

Accurate methods for diagnosis and surveillance of helminth infections are of increasing interest in both human and animal health settings. Such approaches are typically proposed to monitor the presence and ultimately decline of populations targeted by large-scale control measures, such as mass drug administration (MDA) for the prevention and/or treatment of human helminth infections, or prophylactic treatment of domesticated animals. An ideal diagnostic will be sensitive, to detect the parasite if in fact present, and specific, to identify the targeted parasite species in the presence of non-target material such as other parasite species or the host. Ideally, samples taken for diagnostic purposes can be used to gather additional information beyond the presence or absence of a specific parasite, so the same material could be used for example, to predict how well the infection will respond to drug treatment, or how the parasite is related to other endemic or imported parasites. As most parasitic stages of helminths of humans and animals are naturally inaccessible *in vivo* (not accounting for potential availability of some mature stages of helminths following chemo-expulsion, for example, *Ascaris lumbricoides* and *Trichuris trichiura*), a diagnostic should also be informative on non-invasive stages of the parasite, such as eggs deposited in faeces, or intermediate stages of the parasite’s life cycle that exist in the external environment.

A key challenge of working with environmental intermediate stages is that they are often immature, for example, eggs or early stage larvae, and extremely small (for example, *Haemonchus contortus* eggs are approximately 75 × 44 μm and *Schistosoma mansoni* miracidia approximately 140 × 55 μm), limiting the amount of accessible material (e.g., DNA) available to perform the diagnostic assay. They are often environmentally resistant, and the same features that naturally protect the DNA from damage prior to reinfection make it difficult to extract DNA. In many cases they are isolated from host faeces and so are susceptible to bacterial contamination, or from host tissues and so become contaminated with host DNA. Furthermore, samples may need to be transported efficiently to a laboratory setting without a significant loss of this already limited material. A number of approaches have been tested to preserve macromolecules from individual parasites for transport and storage, including ethanol, RNAlater and Whatman® FTA® cards, from which robust PCR and microsatellite data could be profiled (Boué et al., 2017; Campbell et al., 2017; Gower et al., 2007; Marek et al., 2014; Webster, 2009; Webster et al., 2012; Xiao et al., 2013). Although under ideal conditions the detection of a single DNA molecule is possible, the limited material available per parasite has to date largely restricted assaying to a small number of loci, limiting the amount of information obtained from any individual parasite.

Genomic approaches offer an information-rich technology for diagnostic and surveillance applications. Increasing throughput and decreasing costs of whole genome sequencing has resulted in the recent and steadily growing application of genomics in helminth parasitology; for example, for diagnostic applications, high throughput amplicon sequencing for helminth species identification and community composition (Avramenko et al., 2015) and the presence of drug resistance alleles (Avramenko et al., 2019) have been described. Although low DNA concentrations are typically prohibitive for genome-wide approaches on individual parasites, a number of studies have successfully used whole genome amplification on DNA extracted from single larval stages to perform reduced representation (Shortt et al., 2017) and exome (Le Clec’h et al., 2018; Platt et al., 2019) sequencing on miracidia of *Schistosoma* spp., and whole genome sequencing of *Haemonchus contortus* L3 stage larvae (Doyle et al., 2018) and microfilaria of *Wuchereria bancrofti* (Small et al., 2018). Whole genome amplification protocols do, however, add considerable expense per sample, and can introduce technical artefacts such as uneven and/or preferential amplification (potentially of contaminant sequences), chimeric sequences, and allele dropout (Sabina and Leamon, 2015; Tsai et al., 2014), that may lead to a reduction in genetic diversity, and in turn, relevance to the original unamplified material. The field of genomics is, however, rapidly advancing towards very low minimum sample input requirements, and single cell approaches for DNA and RNA sequencing are now available. Such approaches have begun to be used on parasitic species such as *Plasmodium* spp. (Howick et al., 2019; Ngara et al., 2018; Reid et al., 2018; Trevino et al., 2017), but are yet to be adopted by helminth parasitologists. Although these low-input, high-throughput approaches are not designed – and perhaps not currently suitable – for diagnostic applications, the developments in molecular biology techniques for low input sequencing can benefit the use of genomics for helminth applications. Here, we test a number of low-input DNA extraction approaches for individual helminth samples stored on Whatman® FTA® cards, followed by low-input library preparation without whole genome amplification, and whole-genome sequencing. A total of five DNA extraction approaches were initially tested, after which the most promising approach was applied to multiple life stages from eight helminth species. The results presented here demonstrate the advancement of low-input whole genome sequencing, and are discussed in the context of their utility for helminth diagnostics and surveillance.

## Methods

### Sample collection

Samples representing accessible, immature life stages of a total of eight helminth species were tested, the collection of which is described below.

#### Ancylostoma caninum

Fresh feces from a research purpose-bred laboratory beagle (University of Georgia AUP # A2017 10-016-Y1-A0) infected with the Barrow isolate (drug-susceptible isolate from Barrow County Georgia, USA) were collected and made into a slurry with water, filtered through 425 µm and 180 µm sieves, and centrifuged at 2500 rpm for 5 min after which the supernatant was discarded. Kaolin (Sigma-Aldrich, St. Louis, MO) was then added and resuspended in sodium nitrate (SPG 1.25–1.3) (Feca-Med^®^, Vedco, Inc. St Joseph, MO, USA). The tube was then centrifuged at 2500 rpm for 5 min, after which the supernatant was passed through a 30 µm sieve and rinsed with distilled water, and reduced to a volume of 10-15 mL. The volume was adjusted to 1 egg per 5 µL using distilled water. The eggs were stored at room temperature for 2 h before placing them onto the Whatman^®^ FTA^®^ cards. Eggs were also placed onto Nematode growth medium (NGM) plates (Sulston and Hodgkin, 1988) and incubated at 26°C to obtain the first-stage (L1) larvae. After 48 h, larvae were rinsed off the plate with distilled water and centrifuged at 1000 rpm for 5 mins. Larvae were counted and the concentration adjusted to 1 larva per 5 µL. The larvae were stored at room temperature for 2 h before placing them onto the Whatman^®^ FTA^®^ cards. To obtain third-stage (L3) larvae, eggs were isolated from fresh feces from a research purpose-bred laboratory beagle (University of Georgia AUP # A2017 10-016-Y1-A0) infected with the Worthy isolate (Worthy 3.1F3Pyr; multiple-drug resistant isolate originally isolated from a greyhound dog, Florida, USA). Eggs were placed onto NGM plates (Sulston and Hodgkin, 1988) and incubated at 26°C. After seven days, larvae were rinsed off the plate with distilled water and centrifuged at 1000 rpm for 5 mins. Larvae were counted and the concentration adjusted to 1 larva per 5 µL. The larvae were stored at room temperature for 2 h before placing them onto the Whatman^®^ FTA^®^ cards.

#### Ascaridia dissimilis

Eggs of *A. dissimilis* (Isolate Wi: North Carolina, USA) were isolated from excreta of experimentally infected turkeys. Water was added to the excreta and made into a slurry, which was filtered using a 425 µm and 180 µm sieve to remove large debris. The remaining particulates were placed into 50 mL centrifuge tubes and centrifuged at 433 g for 7 mins. Supernatant was removed and the pellet was resuspended in a saturated sucrose solution with a specific gravity of 1.15. The suspension was centrifuged as before and eggs were isolated from the top layer. Eggs were rinsed over a 20 µm sieve with water to remove residual sucrose and then concentrated to 1 egg per 5 µL using deionized water. Multiple 5 µL aliquots of the egg solution were dispensed using a micropipette onto the Whatman^®^ FTA^®^ card.

#### Dirofilaria immitis

Blood was taken from a dog infected with the macrocyclic lactone (ML)-resistant Yazoo strain (Yazoo: originally isolated from a dog in Yazoo City, Mississippi, USA; See Maclean *et al.* (2017) for complete history). To obtain microfilariae, blood was collected in heparin tubes and centrifuged for 30 min at 2500 rpm after which the supernatant was discarded. The pellet was suspended in 3.8% sodium citrate (Sigma-Aldrich, St. Louis, MO) and 15% saponin (Sigma-Aldrich, St. Louis, MO) was added in a 1:7 dilution. The tube was then vortexed and centrifuged for 30 min at 2500 rpm after which the supernatant was discarded and the pellet resuspended in 3.8% sodium citrate to the original blood volume, vortexed, and then centrifuged for 4 min at 2500 rpm. The pellet was then resuspended and mixed in a 1:9 solution of 10⍰ phosphate buffered saline (PBS; Thermo Fisher Scientific, Waltham, MA) and distilled water. The tube was then centrifuged for 4 min at 2500 rpm and the pellet resuspended in PBS. The microfilariae were then counted and adjusted accordingly to have one microfilaria per 5 µL and stored at room temperature for 2 h before placing them onto the Whatman^®^ FTA^®^ cards.

#### Dracunculus medinensis

Individual L1 samples were obtained as progeny of an adult female worm manually extracted from an infected dog in Tarangara village, Chad (9.068611 N, 18.708611 E) in 2016. This extraction forms part of the standard containment and treatment procedure for Guinea worm infections, as agreed upon and sanctioned by the World Health Organisation and country ministries of health. The adult worm was submerged in ethanol in a microcentrifuge tube for storage; L1 stage progeny that were found settled on the bottom of the tube were collected for analysis.

#### Haemonchus contortus

Eggs representing the F5 generation of a genetic cross (described in Doyle *et al.* (2018)) were collected from fresh faeces from experimentally infected sheep housed at the Moredun Research Institute, UK. All experimental procedures were examined and approved by the Moredun Research Institute Experiments and Ethics Committee and were conducted under approved UK Home Office licenses (PPL 60/03899) in accordance with the Animals (Scientific Procedures) Act of 1986. Briefly, faeces were mixed with tap water and passed through a 210 µm sieve, then centrifuged at 2500 rpm for 5 mins in polyallomer tubes. The supernatant was discarded, before adding kaolin to the faecal pellet, vortexing, and resuspending in a saturated salt solution. After centrifugation at 1000 rpm for 10 mins, the polyallomer tube was clamped to isolate eggs, which were collected on a 38 µm sieve and rinsed thoroughly with tap water. Eggs were incubated on NGM plates at 20°C for 48 h to hatch to L1 stage larvae. In addition to freshly collected material, eggs collected in the same manner then stored at −20°C, from a previous generation of the cross, were also tested. Eggs and L1 larvae were resuspended in PBS and spotted onto Whatman FTA cards in 3 µL per egg/L1.

#### Schistosoma mansoni

Three collections of *S. mansoni* samples were used in this work. The first were field samples collected from humans on Lake Victoria fishing villages in Uganda as part of the LaVIISWA trial (Sanya et al., 2018). Ethical approval for this trail was given by the Uganda Virus Research Institute (reference number GC127), Uganda National Council for Science and Technology (reference number HS 1183) and London School of Hygiene & Tropical Medicine (reference number 6187). Parasite eggs were collected from participants’ stool samples using a Pitchford-Visser funnel, washed with mineral water until clean, and transferred into a petri dish with water to be hatched in direct sunlight. After hatching, the miracidia were picked in 2 µL water using a pipette and placed on a Whatman^®^ FTA^®^ card for storage. The second were field samples collected as part of a repeated cross-sectional study of MDA exposure in school children in Uganda. Patient enrolment, including written consent, and sample collection have been described previously (Crellen et al., 2016). Ethical approvals for this study were granted by the Uganda National Council of Science and Technology (MoU sections 1.4, 1.5, 1.6) and the Imperial College Research Ethics Committee (EC NO: 03.36. R&D No: 03/SB/033E). Host stool was sampled 1-3 days prior to treatment with praziquantel (40 mg/kg) and albendazole (400 mg). A Pitchford-Visser funnel was used to wash and filter stool to retain parasite eggs. The filtrate was kept overnight in water and hatched the following morning in sunlight. Individual miracidia were isolated with a 20 μL pipette and transferred into petri-dishes of nuclease-free water twice before spotting onto Whatman^®^ FTA^®^ cards. The third source of miracidia were derived from the livers of experimentally infected mice kept to maintain the *S. mansoni* life cycle at Wellcome Sanger Institute. Mouse infection protocols were approved by the Welfare and Ethical Review Body (AWERB) of the Wellcome Sanger Institute. The AWERB is constituted as required by the UK Animals (Scientific Procedures) Act 1986 Amendment Regulations 2012. BalbC mice (6-8 week old) were infected with 250 cercariae, after which livers were collected on day 40 post infection. Eggs were isolated from the liver tissues using collagenase digestion followed by percoll gradient, and were washed well with sterile PBS, before being hatched in sterile conditioned water. The hatched individual miracidia were spotted onto Whatman^®^ FTA^®^ cards.

#### Strongyloides stercoralis

The *S. stercoralis* UPD strain and the isofemale isolate PVO1 were maintained in purpose-bred, prednisone-treated mix breed dogs according to protocol 804883 approved by the University of Pennsylvania Institutional Animal Care and Use Committee (IACUC), USA. IACUC-approved research protocols and all routine husbandry care of the animals was conducted in strict accordance with the Guide for the Care and Use of Laboratory Animals of the National Institutes of Health, USA. Faeces were collected, moisturized and mixed with equal volume of charcoal and cultured at 22°C in 10 cm plates (Lok, 2007). Post parasitic stage one and two larvae (L1/L2), free-living male and female adults, and infective third-stage (L3) were isolated by the Baermann technique after 24 h, 48 h, and 6 days in these charcoal coprocultures, respectively (Jaleta et al., 2017; Lok, 2007). For this study, free living females, L1 and L3 larvae were collected from the Baermann funnel sediments and washed three times using PBS.

#### Trichuris muris

Infection and maintenance of *T. muris* was conducted as described (Wakelin, 1967). The care and use of mice were in accordance with the UK Home Office regulations (UK Animals Scientific Procedures Act 1986) under the Project license P77E8A062 and were approved by the institutional Animal Welfare and Ethical Review Body. Female SCID mice (6–10 wk old) were orally infected under anaesthesia with isoflurane with a high dose (n = 400) of embryonated eggs from *T. muris* E-isolate. Mice were monitored daily for general condition and weight loss. At day 35 post infection, mice were killed by exsanguination under terminal anesthesia, after which adult worms were harvested from cecums. Adult worms were cultured in RPMI 1640 supplemented with 10% fetal calf serum (v/v), 2 mM L-glutamine, penicillin (100 U/mL), and streptomycin (100 mg/mL; all Invitrogen), for 4 h or overnight, and eggs were collected. The eggs were allowed to embryonate for at least 6 weeks in distilled water, and infectivity was established by worm burden in SCID mice. *T. muris* eggs were hatched to produce sterile L1 larvae using 32% sodium hypochlorite in sterile water for 2 h at 37°C with 5% CO_2_. Eggs were washed with RPMI 1640 supplemented with 10% fetal calf serum (v/v), 2 mM L-glutamine, penicillin (100 U/mL), and streptomycin (100 mg/mL; all Invitrogen), and incubated at 37°C with 5% CO_2_ for 4 to 5 days until they hatched.

For each species, unless otherwise stated, pools of individuals were washed in sterile PBS, before being transferred to a petri dish. Individuals were identified under the microscope, after which 5 µL PBS containing an individual parasite was transferred onto a Whatman^®^ FTA^®^ card and dried for a minimum of 20 mins at room temperature prior to storage or shipping to the Wellcome Sanger Institute, UK. The Whatman^®^ FTA^®^ cards with samples spotted were stored in a clean plastic bag in the dark at room temperature prior to analysis.

### DNA extraction

The sample spots on FTA cards were punched out manually into 96-well plates using either a Harris Punch or autonomously using robotics. The DNA extraction was carried out for each method as described below:

1. *Nexttec* (NXT): Extraction using the Nexttec 1-step DNA Isolation Kit for Tissues & Cells (cat: 10N.904; Waendel Technology Limited, UK) was performed according to manufacturer’s guidelines. 75 µl of proteinase lysis buffer was used for digestion.
2. *Bloodspot* (BSP): The bloodspot extraction was derived from the QIAamp DNA Investigator Kit (cat: 51104/51106; Qiagen), following the “DNA Purification from Dried Blood Spots (QIAamp DNA Mini Kit)” protocol according to manufacturer’s guidelines.
3. *CGP* (CGP): Extraction using the CGP protocol (Moore et al., 2018) involved adding 30 µL of lysis buffer (1.25 µg/mL of Protease reagent (Qiagen; cat# 19155) in Tris HCl pH 8.0, 0.5% Tween 20, 0.5% NP40) to individual Whatman^®^ FTA^®^ punches in a 96-well PCR plate. The samples were incubated at 50°C for an hour followed by protease inactivation by heating the samples to 75°C for 30 min.
4. *ForensicGem* (FGM): Extraction using the ForensicGEM Universal DNA extraction kit (cat: FUN0100; ZyGem) was performed according to the manufacturer’s guidelines.
5. *PicoPure* (PIP): Extraction using the ARCTURUS^®^ PicoPure^®^ DNA Extraction Kit (cat: KIT0103; ThermoFisher) was performed according to the manufacturer’s guidelines, using 75 µL of the extraction buffer.

When the extracted DNA from all the above methods was present in a volume greater than 25 µL, samples were cleaned with Agencourt AMPure XP beads (Beckman-Coulter) and eluted in 25 µL nuclease-free H_2_O. The entire DNA samples were used downstream to make sequencing libraries. A summary of the number of species, life stages, and conditions tested, is presented in **Table 1**.

**Table 1.** Summary of the number of species, life stages, and conditions tested

| Species | Life stage | Extraction kit | Sample number |
| --- | --- | --- | --- |
| *Ancylostoma caninum* | Eggs | CGP | 8 |
| *Ancylostoma caninum* | L1 | CGP | 8 |
| *Ancylostoma caninum* | L3 | CGP | 8 |
| *Ascaridia dissimilis* | Eggs | CGP | 8 |
| *Dirofilaria immitis* | microfilaria | CGP | 8 |
| *Dracunculus medinensis* | L1 | Nexttec | 129 |
| *Haemonchus contortus* | Egg | Bloodspot | 10 |
| *Haemonchus contortus* | Egg | CGP | 10 |
| *Haemonchus contortus* | Egg | ForensicGem | 4 |
| *Haemonchus contortus* | Egg | Nexttec | 10 |
| *Haemonchus contortus* | Egg | Picopure | 10 |
| *Haemonchus contortus* | Frozen Egg | Bloodspot | 6 |
| *Haemonchus contortus* | Frozen Egg | CGP | 6 |
| *Haemonchus contortus* | Frozen Egg | Nexttec | 6 |
| *Haemonchus contortus* | Frozen Egg | Picopure | 6 |
| *Haemonchus contortus* | L1 | Bloodspot | 10 |
| *Haemonchus contortus* | L1 | CGP | 10 |
| *Haemonchus contortus* | L1 | ForensicGem | 4 |
| *Haemonchus contortus* | L1 | Nexttec | 10 |
| *Haemonchus contortus* | L1 | Picopure | 10 |
| *Schistosoma mansoni* | miracidia | Bloodspot | 59 |
| *Schistosoma mansoni* | miracidia | CGP | 36 |
| *Schistosoma mansoni* | miracidia | ForensicGEM | 10 |
| *Schistosoma mansoni* | miracidia | Nexttec | 51 |
| *Schistosoma mansoni* | miracidia | PicoPure | 12 |
| *Strongyloides stercoralis* | FL | CGP | 8 |
| *Strongyloides stercoralis* | iL3 | CGP | 8 |
| *Strongyloides stercoralis* | L1 | CGP | 8 |
| *Trichuris muris* | bleached eggs | CGP | 8 |
| *Trichuris muris* | eggs | CGP | 8 |
| *Trichuris muris* | L1 | CGP | 8 |

### Library preparation and Sequencing

DNA sequencing libraries for all samples were prepared using a protocol designed for library preparation of laser capture microdissected biopsy (LCMB) samples using the Ultra II FS enzyme (New England Biolabs) for DNA fragmentation as previously described (Lee-Six et al., 2019). A total of 12 cycles of PCR were used (unless otherwise stated in **Table S2**) to amplify libraries and to add a unique 8-base index sequence for sample multiplexing.

Multiplexed libraries were sequenced using the Illumina MiSeq system with V2 chemistry 150 bp paired end (PE) reads. The *D. medinensis* samples were sequenced as part of a different study using the HiSeq 2500 with V4 chemistry 125 bp PE reads. In general, we performed sufficient low coverage sequencing on each sample to enable us to identify: (i) the proportion of on-target mapped reads, (ii) the proportion of duplicate reads, i.e. library artefacts, and (iii) the proportion of off-target contaminant reads.

Metadata for each sample, including sample IDs, sequencing lane IDs, ENA sample accession numbers, and data generated are described in **Table S2**. Raw sequence data will be made available under ENA study ID ERP114942.

### Analysis

Reference genomes from each of the test species were obtained from WormBase Parasite (Howe et al., 2017) Release 12. Raw sequence data for each species were mapped to their respective reference genome using BWA-MEM (Li, 2013), after which duplicate reads were marked using Picard (v2.5.0; https://github.com/broadinstitute/picard). Samtools flagstats and bamtools stats were used to characterise the outcome of the mapping, the results of which were collated using MultiQC (Ewels et al., 2016). Data were manipulated and visualised in the R (v3.5.0) environment using the following packages: ggplot2 (https://ggplot2.tidyverse.org/), patchwork (https://github.com/thomasp85/patchwork), and dplyr (https://dplyr.tidyverse.org/).

The code to reproduce the analysis and figures for this manuscript is described in https://github.com/stephenrdoyle/helminth_extraction_wgs_test.

## Results & Discussion

The aim of this work was to determine the feasibility and efficiency of using a low input DNA extraction and library preparation approach for whole-genome sequencing of individual helminth parasites. We have targeted immature life stages that are found in the environment for which it is possible to collect samples non-invasively. We first tested our approaches on the nematode *H. contortus* and the trematode *S. mansoni*. Five approaches were tested using single egg (fresh and frozen) and L3 of *H. contortus*, and miracidia of *S. mansoni.* We determined the success of the library preparation approach by comparing the proportion of reads mapped (**Figure 1 & 2**; top plots), representing on-target mapping as a measure of specificity, as well as the proportion of duplicate reads (**Figure 1 & 2**; bottom plots), which typically represent library preparation artefacts due to over-amplification of DNA during PCR. Finally, we determined the proportion of reads associated with off-target contamination by comparison of raw reads to a kraken contamination database (**Table S2**; kraken_unassigned).

**Figure 1.**
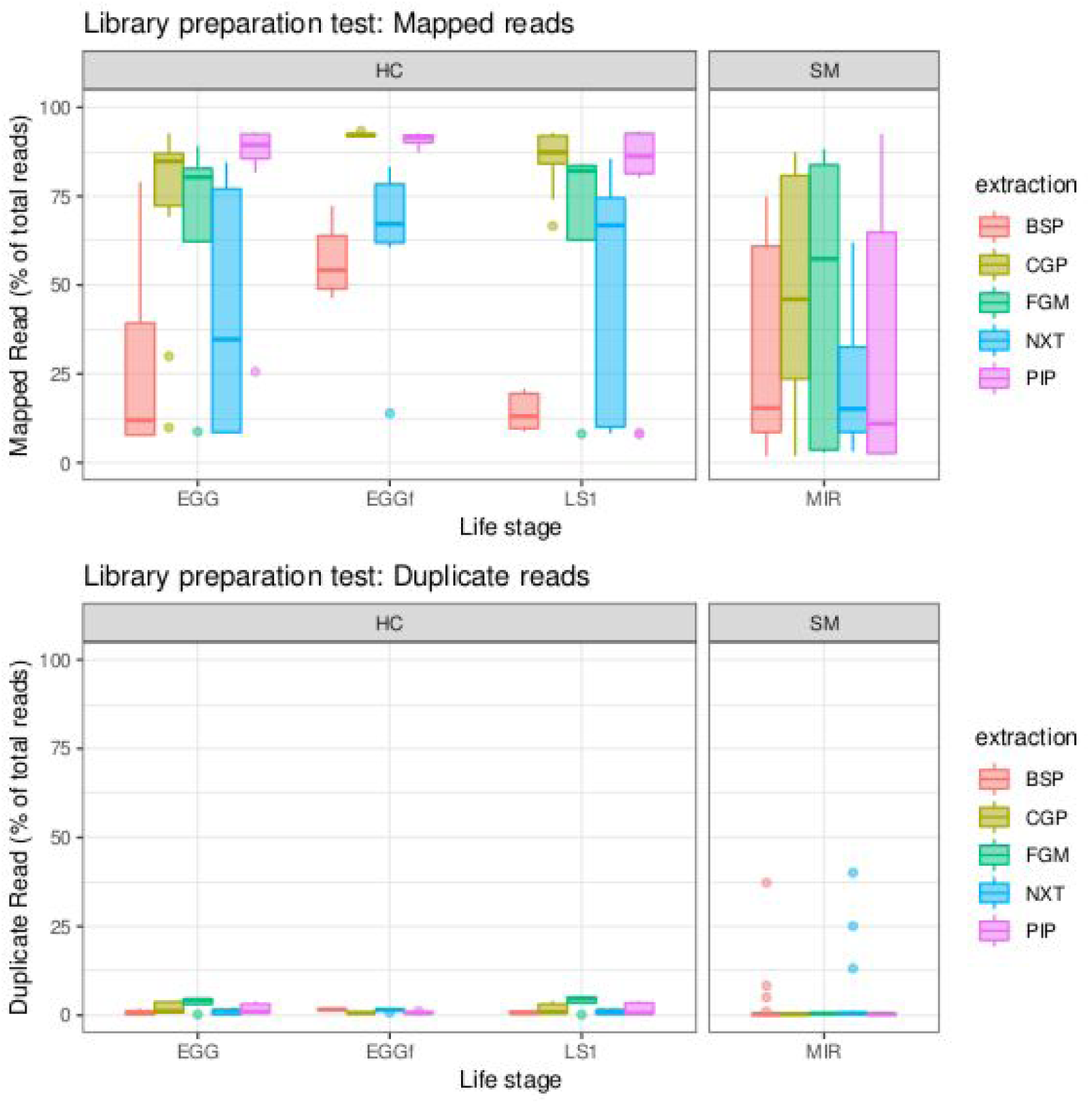
Comparison of library preparation approaches on *Haemonchus contortus* eggs (EGG), frozen eggs (EGGf), and L1 (LS1) stages, and *Schistosoma mansoni* miracidia (MIR). The top plot presents the percentage of reads that mapped to each reference genome, and the bottom plot presents the proportion of duplicate reads identified. Boxplots are coloured by the library preparation approach used. The number of samples in each comparison is presented in Table 1.

There were distinct differences in overall mapping between *H. contortus* and *S. mansoni*; while some of this variation may reflect differences in extracting DNA from different species, the *S. mansoni* samples generally had a greater proportion of contaminating DNA present relative to the *H. contortus* samples (**Table S2**; kraken_unassigned). We achieved some on-target mapping under all optimisation conditions tested, however, significant variation was observed between approaches (**Figure 1**). BSP and NXT generally performed poorly, with either low (median = 19.42; median absolute deviation (MAD) = 16.50) or significant variance (median = 66.20; MAD = 26.18) in mapping frequency between samples observed, respectively. PIP performed consistently well with high mapping rates across all stages in *H. contortus* (median = 90.30; MAD = 3.96), however, it was poor in *S. mansoni* (median = 11.00; MAD = 12.73). The duplication rate of all conditions were within an acceptable low range (median = 0.46, MAD = 0.4), with only 2% of samples having greater than 5% duplicate reads. CGP and FGM performed most consistently between stages and species; FGM had higher variance and duplication rates relative to CGP across all samples tested, and therefore, CGP was chosen to explore further.

We expanded our analysis to a total of eight helminth species for which samples were available, including a total of six distinct life stages (**Figure 2**). All tested samples were extracted with the CGP protocol, except for *D. medinensis*, which were extracted using NXT and included for comparison. High variability in mapping was observed between species, with 50% or greater mapping frequency achieved in at least one life stage of 6 of the 8 species tested. Clear differences were observed between multiple life stages tested within a species, likely reflecting differences in extraction efficiency, for example: for *A. canium,* reads from eggs (median = 54.98) mapped much more effectively than L1 (median = 3.61) or L3 stages (median = 7.57), and in *S. stercoralis,* full-length females (median = 67.84) and L1 (median = 47.19) performed better than L3 (median = 9.01). Interestingly, L1 larvae (median = 41.85) and bleached eggs (median = 51.35) performed much better than untreated eggs (median = 2.72) of *T. muris*; bleaching is experimentally used for promoting hatching of *T. muris*, by dissolving the egg shell layers and in turn improving access to DNA within. However, bleached eggs were embryonated and developmentally more advanced than untreated unembryonated eggs, and therefore would have more DNA available for library preparation. Similar to untreated *T. muris* eggs, *A. dissimilis* eggs performed poorly (median = 0.16); for both species, few if any nematode sequences were recovered and the majority of sequencing reads were contaminating bacterial-derived contaminants, perhaps indicative of the challenge of accessing material with the environmentally-resistant egg. Analysis of *A. dissimilis* was further limited by the lack of a reference genome for this species; we mapped against the available genome of *Ascaris lumbricoides*, the nearest species for which a reference was available, and therefore at best, would have expected suboptimal mapping due to sequence divergence. The duplication rates remain low for all species and conditions tested, apart from *D. medinensis*, which was much higher (median = 36.90). This was unexpected, given NXT did not produce high duplication rates under the initial conditions tested with *H. contortus* or *S. mansoni*, nor were there excessive PCR cycles used in this instance.

**Figure 2.**
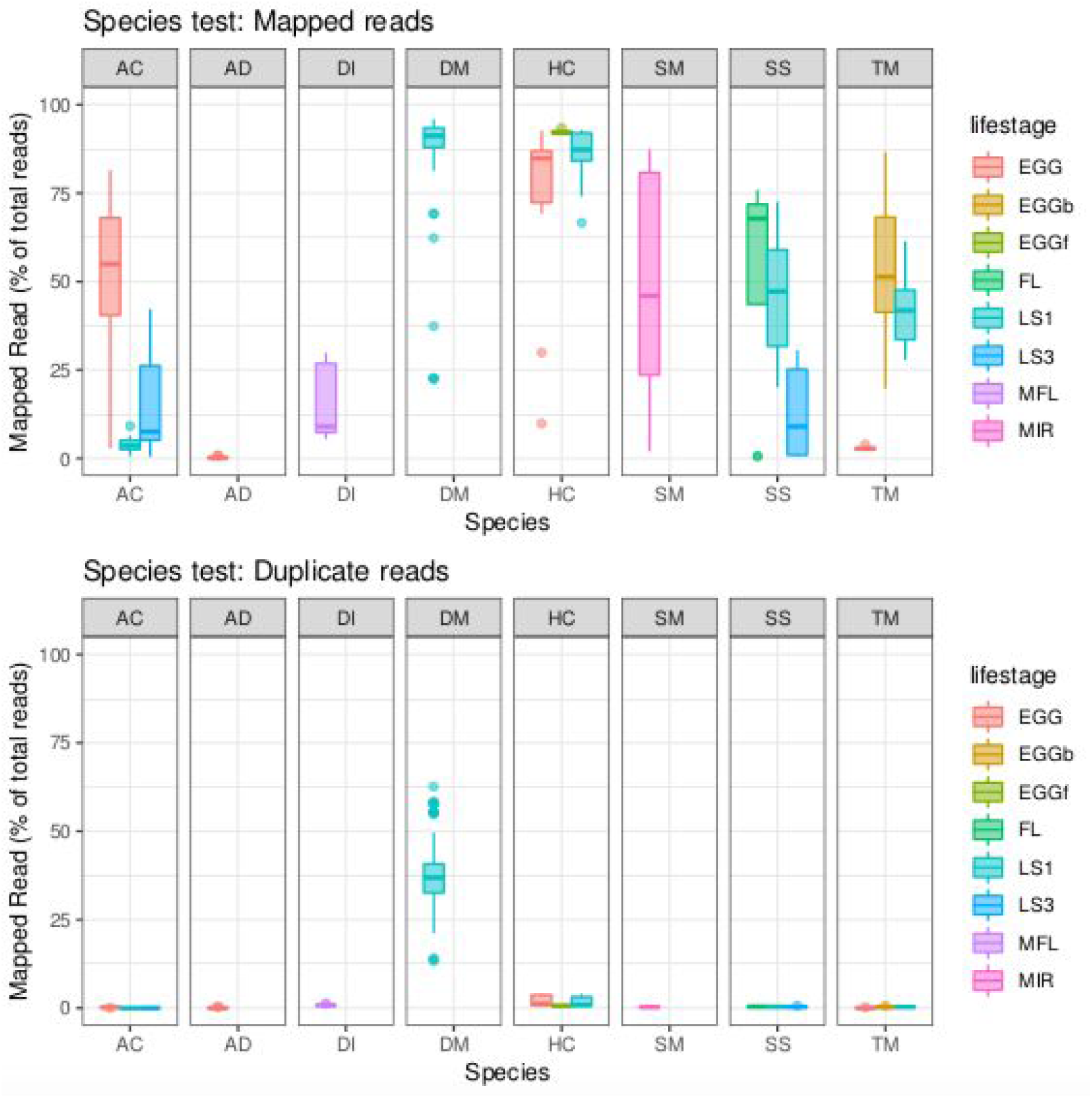
Comparison of library preparation from multiple life stages of 8 helminth species. All libraries were prepared with the CGP approach, except for *D. medinensis*, which was prepared with NXT and presented alongside for comparison. As in Figure 1, the top plot presents the percentage of reads that mapped to each reference genome, and the bottom plot presents the proportion of duplicate reads identified. Boxplots are coloured by the life stage assayed. Egg = untreated egg; EGGb = bleached egg; EGGf = frozen egg; FL = full-length female; LS1 = larval stage 1; LS3 = larval stage 3; MFL = microfilaria; MIR = miracidia. The number of samples in each comparison is presented in Table 1.

In summary, we present successful DNA extraction followed by whole-genome sequencing of individual parasites from 6 of 8 species examined. These results significantly extend the possibility of genomic analyses for life stages for which, at best, were limited to low-resolution, low-throughput PCR based assays without the addition of whole genome amplification. Whatman^®^ FTA^®^ cards provide a convenient substrate for sample collection and storage, and do not limit the application of direct DNA extraction and whole genome sequencing of parasite samples, even for field samples as demonstrated for *S. mansoni* miracidia that were collected and processed in Uganda before they were transported to the UK. Further optimisation is required to improve the DNA recovery from eggs, for example, from *A. dissimilis* and *T. muris*, to provide greater applicability of our approaches to species that generate particularly environmentally-resistant stages, such as the soil transmitted helminths. The application of whole genome sequencing to diagnose and monitor helminth infections at scale is largely limited by the costs of library preparation and sequencing, and therefore, will be restricted to niche applications of the technology. However, targeting the mitochondrial genome, for example, by whole genome sequencing may be a viable and cost-effective alternative, potentially providing greater diagnostic information than low throughput PCR-based diagnostics (**Table 2**). Continued development of genomic technologies and the associated reduction in sequencing and library preparation costs will make screening large samples by genome sequencing more routine as in viral (Dudas et al., 2017) and bacterial (Domman et al., 2018) population studies. In doing so, the ability to derive information-rich data for diagnostic and surveillance purposes using genomics (Cotton et al., 2018) will be particularly informative as efforts to control human infective helminths using MDA move from control to elimination.

**Table 2.** Breakdown of sequencing strategies per species based on whole genome sequencing at 30X coverage and whole genome sequencing to achieve 100X whole mitochondrial genome coverage

| Species / stage | Genome assembly size <sup>1</sup> | Number of samples that can be multiplexed <sup>2</sup> | mtDNA/ nuclear ratio <sup>3</sup> | Sampled multiplexed targeting mtDNA at 100X coverage <sup>2</sup> | Nuclear genome coverage per 100X mtDNA genome |
| --- | --- | --- | --- | --- | --- |
| *S. mansoni* miracidia | 409 | 173 | 142.77 | 7418 | 0.70 |
| *H. contortus* egg | 283 | 250 | 70.76 | 5313 | 1.41 |
| *H. contortus* L1 | 283 | 250 | 44.55 | 3345 | 2.24 |
| *D. medinensis* L1 | 103 | 688 | 70.81 | 14608 | 1.41 |
| *D. immitis* microfilaria | 88 | 805 | 14.27 | 3446 | 7.01 |
| *S. stercoralis* FL | 42 | 1687 | 11.06 | 5597 | 9.04 |

1. Genome size: Obtained from Wormbase Parasite v12 2. Multiplex: total number of samples per NovaSeq 6000 sequencing run with S4 2x150 bp PE chemistry generating 2.5 Tb of data and accounting for 85% mapping rate. Theoretically possible, but may be limited by barcoding available. 3. mtDNA/nuclear ratio: based on coverage of mitochondrial and nuclear derived sequencing reads, normalised to the nuclear genome coverage

## Acknowledgments

Work performed at the Wellcome Sanger Institute is supported by Wellcome Trust (grant 206194) and by the Biotechnology and Biological Sciences Research Council (BB/M003949/1). We thank Alison Elliot for access to *S. mansoni* samples collected from Uganda, the collection of which was supported by Wellcome Trust (grant 095778/Z/11/Z). *S. mansoni* samples were also obtained from the Schistosomiasis Collection at the Natural History Muesem (NHM) [SCAN], which is funded with support from the Wellcome Trust (grant no. 104958/Z/14/Z). We thank the Carter Center for supporting molecular work on *D. medinensis,* and the Guinea worm eradication program for making samples available.

## Author contributions

Conceptualization: SRD, JAC, NH

Methodology: SRD, GS, NH

Software: SRD

Formal analysis: SRD

Investigation: SD, GS

Resources: FA, TJ, JL, JBC, PJC, JPW, TC, ET, TC, MAD-C, PE, RL, KM, CM, TM, BS, PTO, JVT, AW, ED, RK, DB, MB

Data Curation: SRD, NH

Writing – original draft preparation: SRD

Writing – review and editing: All authors

Visualisation: SRD

Supervision: SRD, JAC, NH

Project administration: NH

## Supplementary Data

Table S1 - Summary of total samples analysed by species, life stage, and DNA extraction protocol (excel workbook)

Table S2 - Complete metadata per sample (excel workbook)

